# Intratumoral Delivery of Chlorine Dioxide Exploits its ROS-like Properties: A Novel Paradigm for Effective Cancer Therapy

**DOI:** 10.1101/2023.11.24.568512

**Authors:** Xuewu Liu, Zhaoyang Liu, Xueyan Liu, Shuangning Liu, Jiao Zhang

## Abstract

Our study underscores the potential of chlorine dioxide, resembling reactive oxygen species, in promoting tissue regeneration and executing similar functions. Direct intratumoral administration of chlorine dioxide conveniently and sustainably kills cancer cells, without the risk of developing drug resistance. Given that tumors are viewed as non-healing wounds, the regenerative capabilities of chlorine dioxide offer an innovative method to expedite wound healing and enhance patient outcomes in cancer treatment. Compared to traditional methods, chlorine dioxide treatment reduces complications and risks, presenting a safer alternative. Notably, higher concentrations of chlorine dioxide in promoting tumor ablation can particularly demonstrate the potential to disrupt tumor vasculature. We advocate for the intratumoral injection of chlorine dioxide to achieve direct tumor ablation and boost antitumor immunity while minimizing harm to healthy tissues. Although initial findings indicate that chlorine dioxide stimulates an antitumor immune response, further investigation is essential. As oncology advances with the use of immune checkpoint inhibitors, intratumoral chlorine dioxide administration emerges as a promising strategy, potentially extending survival and alleviating treatment burdens. Future research should focus on refining chlorine dioxide protocols, exploring synergistic therapies, and understanding its systemic immune effects, thereby establishing chlorine dioxide’s role in cancer therapy.

**Highlights:**

- Chlorine dioxide demonstrates potent ROS-like capabilities for both cancer elimination and tissue regeneration, offering a and effective treatment approach.
- Direct intratumoral administration of chlorine dioxide conveniently and sustainably kills cancer cells, without the risk of developing drug resistance.
- Chlorine dioxide can enhance tumor ablation potential by disrupting tumor vasculature, contributing to its overall effectiveness.
- Chlorine dioxide promotes an immune response against cancer, significantly enhancing its therapeutic effectiveness.
- Intratumoral injections of chlorine dioxide represent an innovative and promising strategy for efficacious cancer therapy.

## Introduction

Certain natural compounds or drugs that target Superoxide dismutase (SOD) possess a unique capability to selectively eradicate cancer cells by stimulating the generation or accumulation of Reactive Oxygen Species (ROS) [1, 2]. Furthermore, the body’s neutrophils can also generate ROS, which contributes to the destruction of cancer cells [3]. Certain drugs can selectively enhance the levels of ROS in cancer cells, exhibiting long-term inhibitory effects on cancer. For example, the combination of metformin-induced ROS upregulation and apigenin amplification results in significant anticancer activity while preserving the integrity of normal cells [4]. Photodynamic therapy (PDT) is a treatment method that utilizes light to activate a photosensitizer within the body, resulting in the production of singlet oxygen (^1^O_2_). This sudden increase in oxygen levels induces toxicity in tumor cells, leading to their demise through apoptosis or necrosis. PDT not only directly targets and eradicates the tumor but also stimulates the release of cell death-associated molecular patterns (DAMPs) by dendritic cells, activating the immune system’s antigen-presenting response and promoting an anti-tumor immune response [5].

Based on our hypothesis, we propose that the exogenous supplementation of ROS or their analogs could effectively eradicate tumors and, similar to PDT, elicit an antitumor immune response. In this study, we have selected chlorine dioxide (CD) as a ROS-like oxidant to evaluate its potential in cancer therapy. CD is widely acknowledged as a potent oxidizing agent with disinfectant properties, distinguished by its molecular formula ClO_2_. Its applications encompass various domains, including water treatment, food processing, medical hygiene, and environmental cleaning. Notably, CD exhibits rapid destruction of cell membranes and DNA in bacteria, viruses, and other microorganisms, thereby effectively eliminating pathogens and contaminants. Remarkably, at lower concentrations of 0.25 mg/L, CD can eradicate 99% of E. coli (15,000 cells/mL) within a mere 15 seconds [6]. Furthermore, studies have demonstrated that CD and hydrogen peroxide demonstrate comparable efficacy in inducing cell death in human gingival fibroblasts [7].

ROS are generated during cellular respiration for ATP production [8]. However, the dispersed nature of ROS production in the body and the limited capacity to rapidly eliminate a large number of cells, especially within tumor tissues, present significant challenges. These challenges are primarily attributed to tumor hypoxia, characterized by low oxygen levels in the tumor microenvironment [9]. To establish a novel paradigm for cancer treatment, it is crucial to elevate the concentration of CD to a level that can effectively eradicate substantial tumor masses. This concentration exceeds the typical endogenous ROS production in the body. Furthermore, considering the short half-life of CD as an oxidizing agent in body tissues and the imperative of minimizing systemic side effects, intratumoral administration of CD has been selected as the preferred approach to achieve our objective.

## Materials and methods

### Cell Culture

An MTT assay was conducted to assess the inhibitory effect of a CD solution on the survival rate of MCF-7 breast cancer cells, human cardiac myocytes (HCM), and human vascular endothelial cells (HUVECs). Additionally, the MTT assay was utilized to determine the survival inhibition rate of ten non-small cell lung cancer (NSCLC) cell lines, namely H2110, H1975, H650, H1623, H2126, HCC827, A549, H810, H1048, and H1355. To evaluate apoptosis or necrosis in MCF-7 breast cancer cells and human cardiac myocytes, flow cytometry with Annexin-V PI double staining was employed. The cells were incubated with varying concentrations of CD for 24 hours at 37°C, harvested, and washed with DMEM. Subsequently, the cells were resuspended in annexin V-FITC and PI staining solution, followed by a 15-minute incubation in the dark at room temperature. After the addition of binding buffer, the stained cells were analyzed using a FACSCalibur flow cytometer, with FITC fluorescence measured between 515 and 545 nm and PI fluorescence measured between 564 and 606 nm.

### Reagent

CD solution at concentrations of 7.5 mg/mL, 8mg/mL, 13mg/mL, and 15mg/mL was obtained from Beijing Wanbincell Biotechnology Co., Ltd.

### Animal Models

C57BL/6 and Balb/c mice were obtained from the China Experimental Center for Food, Drugs, and Biological Products. All experiments were conducted with the approval of the The Cancer Institute and Hospital, Chinese Academy of Medical Sciences.

### Mouse Safety Studies

C57BL/6 mice were administered a single intraperitoneal injection of 0.3 mL of a CD solution (7.5 mg/mL), which was determined to be well-tolerated. Similarly, a single dose of 0.5 mL of a CD solution (1 mg/mL) was found to be safe for C57BL/6 mice. On the other hand, intracranial injection of a single dose of 0.02 mL of a CD solution (1.5 mg/mL) was deemed safe for C57BL/6 mice.

### CD Simulates the Impact of ROS on Healthy Tissues

Two groups of female C57BL/6 mice, aged 9-10 weeks, were randomly assigned into two groups, with each group consisting of four mice (n=4). The first group received a subcutaneous injection of 0.3 mL of a CD solution with a concentration of 7.5 mg/mL, while the second group received a subcutaneous injection of 0.3 mL of a CD solution with a concentration of 15 mg/mL. The injection sites were monitored daily, and the extent of injury was assessed and recorded using an injury score. The injury score was determined by multiplying the area of the lesion by the severity of the lesion.

### Effect of CD on Tissue Regeneration in Mice

Twenty female C57BL/6 mice (9-10 weeks old) were anesthetized with halothane and had their tails severed 2 cm from the base. The mice were randomly divided into 5 groups, each consisting of 4 mice. Group 1 immersed their tail wounds in a 15mg/mL CD solution for 1 minute and received daily treatment with the same solution. Group 2 immersed their tail wounds in physiological saline solution for 1 minute and received daily treatment with the same solution. Group 3 immersed their tail wounds in a 15mg/mL CD solution for 10 minutes daily for 8 consecutive days, and then on days 10, 14, and 18. Group 4 immersed their tail wounds in a 1.5% hydrogen peroxide solution for 10 minutes daily for 8 consecutive days, and then on days 10, 14, and 18. The progress of wound healing and tail severance wound scores were recorded. Group 5 immersed their tail wounds in a 15mg/mL CD solution for 10 minutes on days 1, 4, 8, 12, 16, and 20. Wound healing progress was assessed using a scale ranging from 0 (fully healed) to 90 (exudate or crusting area).

### Effects of CD Injection in Mouse Tail Vein

Eight female C57BL/6 mice (9-10 weeks old) received a single tail vein injection of 0.2 mL of a CD solution (7.5 mg/mL), with the injection site located 1 cm from the base of the tail. The mice were observed daily for the first 10 days, and photographs were taken on days 10, 30, 50, and 70 for record-keeping.

### Inhibition of Lewis Lung Cancer in Mice by Intrapericardial Injection of CD Solution

A total of ten male C57BL/6 mice, aged 9-10 weeks, were included in the study. The mice were subcutaneously inoculated with 4×10^6^ LLC cells in the right axillary region and subsequently divided into two groups, with ten mice in each group. Starting from the 4th day after cell inoculation, the CD solution was intratumorally injected at a dosage of 0.2mL per mouse (8 mg/mL) every other day for a total of three doses. From the second injection onwards, the dosage was increased to 0.3mL per mouse. The control group received intratumoral injections of PBS injection solution. The experimental period lasted for 10 days, after which all animals were humanely euthanized two days following the administration of the final dose in the experimental group. Tumor weight was measured after recording the animals’ body weight, and the tumor suppression rate was calculated. Tumor length and width were regularly measured to accurately calculate tumor volume.

### Inhibition of B16 Melanoma Growth in C57BL/6 Mice by Intratumoral Injection of CD Solution

Each male C57BL/6 mouse, aged 9-10 weeks, was subcutaneously inoculated with 2×10^6^ cells in the right axillary region. Subsequently, the mice were randomly divided into three groups, with ten mice in each group. On the 8th day after the inoculation of B16 cancer cells, the intratumoral injection group received CD solution intratumorally. The CD solution was administered at a dosage of 0.2mL (13mg/mL) every 4 days for a total of three doses. On the 2nd day after the inoculation of melanoma, the mouse tail vein injection group received CD solution via the tail vein. The CD solution was injected at a dosage of 0.2mL (1.5 mg/mL) every 4 days for a total of five doses. The tumor model group served as the control and received intratumoral injections of PBS injection solution for observation purposes. The experiment was conducted over a period of 20 days. Four animals from each group were humanely euthanized four days after the last dose, and their tumor weight was measured after recording their body weight in order to calculate the tumor suppression rate. On the 20th day, the remaining mice in each group were tested for cytokines.

### Inhibition of Melanoma B16 Metastasis in C57BL/6 Mice by CD Solution

Each male C57BL/6 mouse, aged 9-10 weeks, was subcutaneously inoculated with 2×10^6^ B16 cells in the right axillary region and 1×10^6^ B16 cells in the tail vein. Subsequently, the mice were randomly divided into three groups, with 9 mice in each group. The groups consisted of a tumor model control group, which received intratumoral injections of PBS solution; an intratumoral administration group in the axillary region; and an intratumoral administration group in the axillary region with daily 2-hour inhalation treatment for the tumor-bearing mice.

The administration of CD solution began on the 7th day after melanoma inoculation. The intratumoral injections of CD solution were administered at a volume of 0.2mL (15 mg/mL) every 4 days for a total of 4 doses. In the inhalation group, the mice received intratumoral injections of CD solution at the same dosage and frequency, along with 2 hours of daily inhalation treatment. For the inhalation treatment, 20mL of 15 mg/mL CD solution was placed in the mouse cage for natural evaporation and inhalation.

On the 17th day, three mice from each group underwent HE staining of the tumor tissue and lungs. On the 21st day, the lungs were harvested, and the pulmonary metastatic foci were counted.

### Inhibitory Effect of Intratumoral CD Solution on 4T1 Breast Cancer Cell Transplantation in Balb/c Mice

A total of 32 female Balb/c mice, aged 9-10 weeks, were utilized for this experiment. They were divided into four groups, each consisting of eight mice. The large tumor group (LT group) received a subcutaneous inoculation of 4×10^5^ 4T1 cells in the right axillary region and contralateral mammary pad. Subsequently, they were randomly assigned to two groups: the control group (PBS+LT) and the treatment group (CD+LT), with eight mice in each group. Similarly, the small tumor group (ST group) received a subcutaneous inoculation of 2×10^5^ 4T1 cells in the right axillary region and contralateral mammary pad. They were then divided into the control group (PBS+ST) and the treatment group (CD+ST), with eight mice in each group.

Starting from the 15th day after inoculation, groups 2 and 4 (CD+LT and CD+ST) received intratumoral injections of a CD solution with a concentration of 15 mg/mL. A total volume of 0.2 mL (15 mg/mL) was administered every 3 days for a total of 6 doses. The mammary pad tumors in the control groups (PBS+LT and PBS+ST) were observed without any treatment (Untreated). On the 27th day, partial immune cells were tested, and on the 33rd day, the animals were euthanized. The lungs were harvested for counting pulmonary metastasis, as well as for the detection of serum and axillary tumor cytokines.

On the 27th day of the 4T1 model, blood and spleen samples were collected from 3 mice in each group, totaling 12 mice. Whole blood samples were collected from 12 mice, and spleen samples were collected from the remaining 12 mice. The spleen suspended cells were prepared for the detection of CD3, CD4, CD8, and CD335 using four-color staining, as well as CD11b, Gr-1, and F4/80 using three-color staining. Fresh anticoagulated peripheral blood from the mice (spleen suspended cells) was added to the bottom of the flow cytometry sample tube, and an appropriate amount of fluorescently labeled antibodies was added, including CD3e PerCP-Cy5.5, CD4 PE, CD8a FITC, and CD335 (NKp46) APC (Biolegend), as well as Gr-1 FITC, F4/80 PE, and CD11b APC (Biolegend). Analysis was performed using a flow cytometer (BD).

On the 33rd day of the 4T1 model, blood and axillary tumor samples were collected from 5 mice in each group, totaling 20 mice (Untreated). ELISA was used to measure the levels of IL-6, TNF-α, INF-γ, and CTLA-4 in the serum and tumor samples.

### Statistics

Statistical analyses were performed using Origin 9.0 software. When the transformed data showed significant variation among treatments, differences among populations and treatments were assessed using the nonparametric Mann-Whitney test. However, in most cases, no significant variation was observed among treatments. In such instances, two-tailed Student’s t-tests were used to evaluate differences between two treatments. For comparisons involving more than two treatments, a two-way ANOVA followed by Dunnett’s multiple comparisons tests was employed to compare multiple groups with repeated measures.

## Results

### In Vitro or Normal Tissue Injection Experiments with CD

The initial safety experiments confirmed the safe administration of CD solution at various concentrations to the mice. CD exhibited cytotoxicity against both cancer cells and normal cells, including cardiomyocytes and vascular endothelial cells (Fig. 1A, B). It induced cell death via apoptotic and necrotic pathways, regardless of the cell type (Fig. 1C, D).

**Fig. 1.**
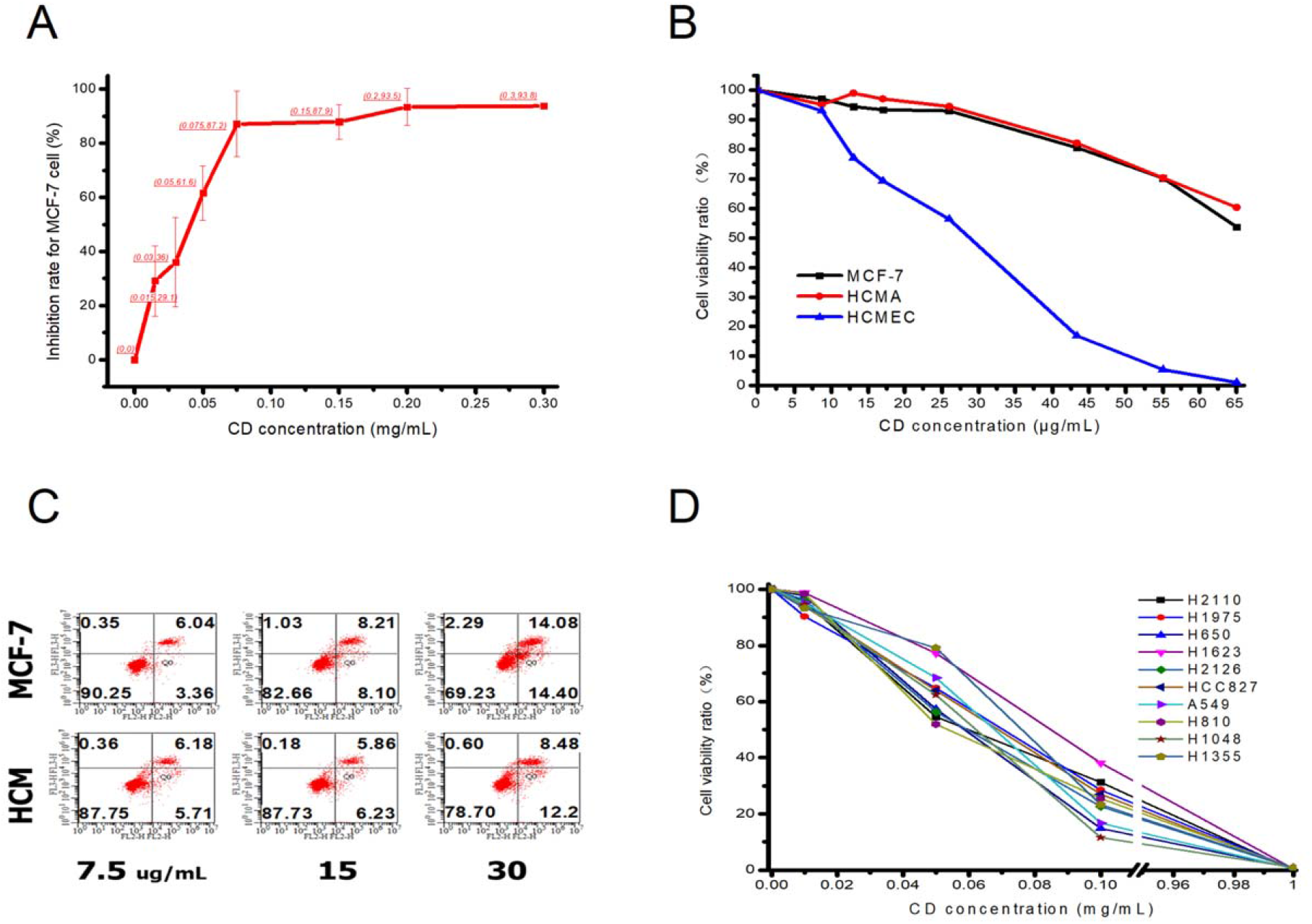
In vitro cell viability assay of CD. (A) Inhibition of cell survival in MCF-7 human breast cancer cells by CD. (B) Percentage of cell survival inhibition by CD in MCF-7 breast cancer cells, normal human cardiomyocytes (HCM), and human vascular endothelial cells (HCMEA). (C) Representative plots demonstrating the measurement of apoptosis or necrotic pathways in breast cancer cells and cardiomyocytes using flow cytometry after treatment with different concentrations of CD (The lower right quadrant represents early apoptotic cells, and the upper right quadrant represents necrotic or late apoptotic cells). (D) Measurement of cell proliferation in 10 NSCLC cell lines when treated with increasing doses of CD.

Subcutaneously injecting 0.3 mL of CD (15 mg/mL and 7.5 mg/mL) into the dorsal region of mice resulted in significant tissue damage, including skin damage and necrosis of various subcutaneous cell types. Higher concentrations of CD caused more severe wounds, but there was no difference in wound healing time. Approximately 20 days later, the damaged tissue completely regenerated, and the skin returned to its original state with successful hair regrowth (Fig. 2A, B; Supplementary Fig. S1D).

**Fig. 2.**
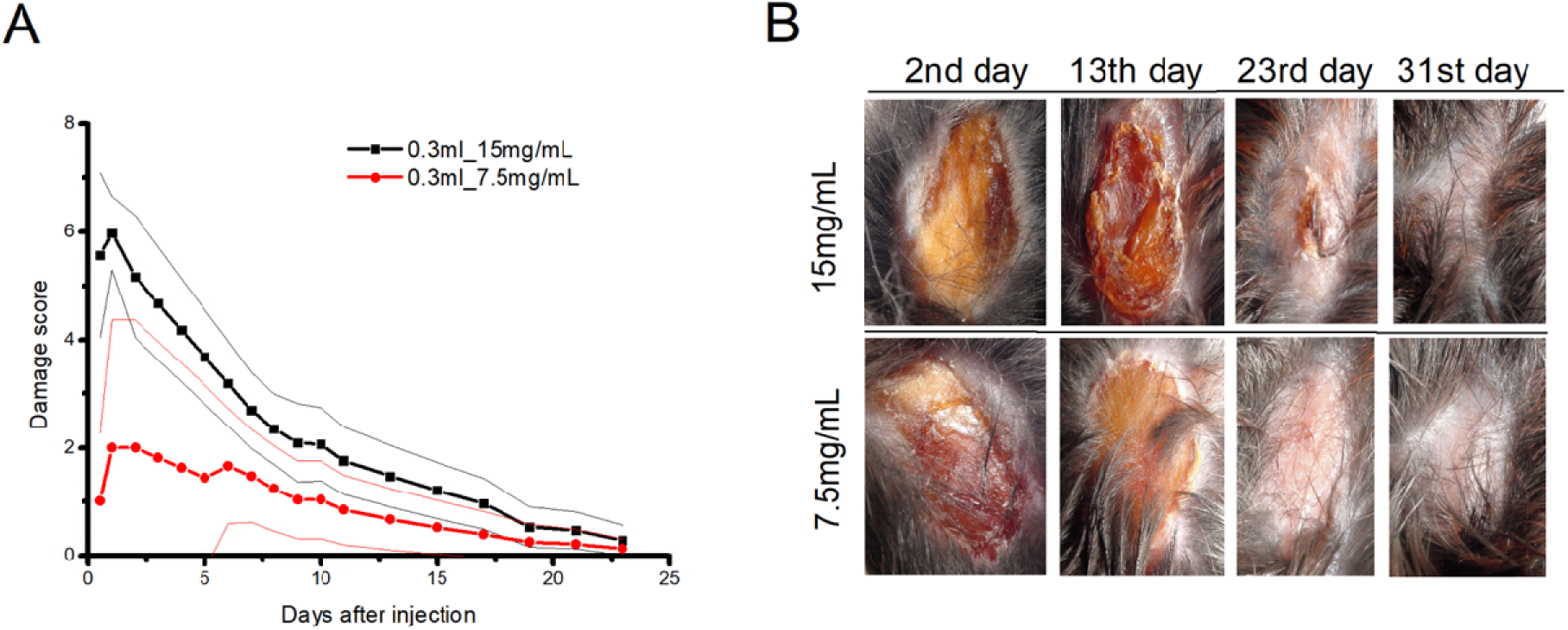
In vivo tissue damage and regeneration assay of CD. (A) Subcutaneously inject 0.3 mL of CD solution at concentrations of 15 mg/mL and 7.5 mg/mL into the dorsal region of C57BL/6 mice, showing the trend of injury scores (injury recovery) over time. Data are presented as mean ± SD (n=4). (B) Representative images of the back skin taken on days 2, 13, and 23 after the start of CD injection. Images selected from each group.

In the tail vein injection group, about 7 to 10 days after injection, mice that received 0.2 mL of CD solution (7.5 mg/mL) showed tail loss of approximately 1-2 cm at the injection site. After 70 days, ischemic necrosis developed in the remaining tail segment distal to the injection site, which eventually fell off (Fig. 3).

**Fig. 3.**
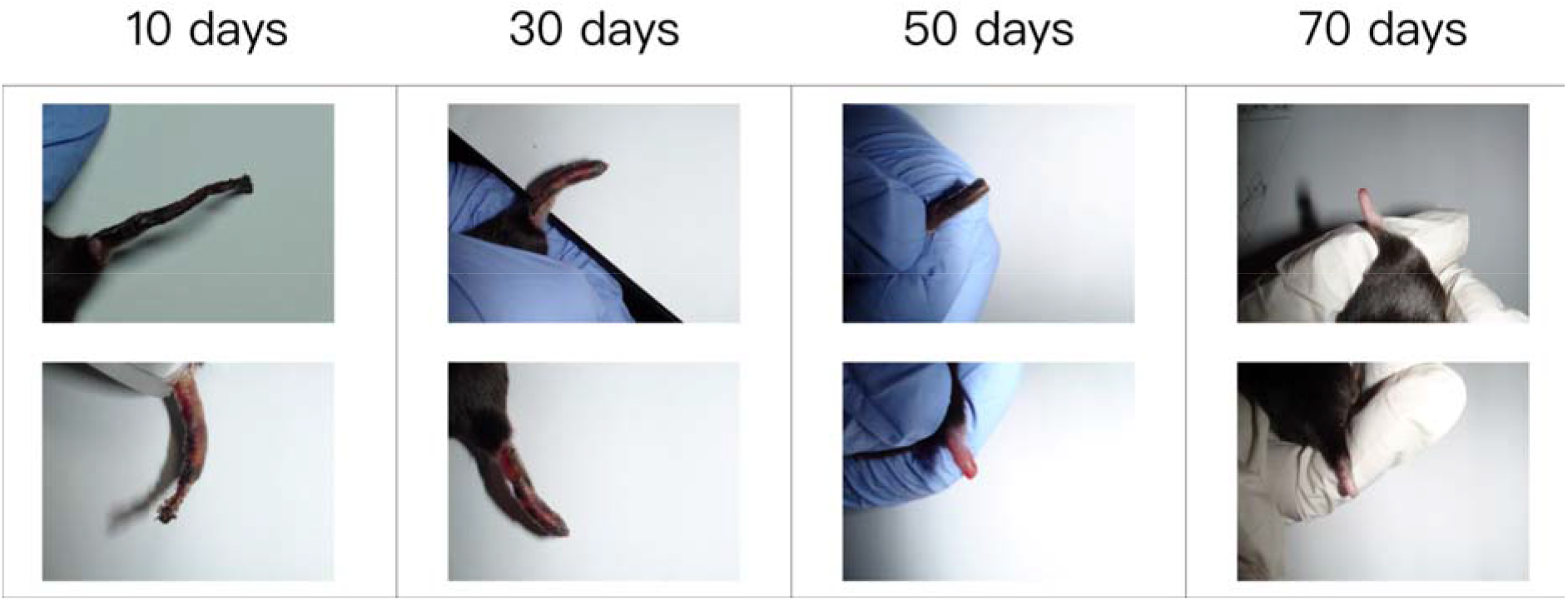
In the tail vein injection experiment with CD. We randomly selected photos of the tails of two mice at time points: 10 days, 30 days, 50 days, and 70 days after the injection of 0.2 mL of CD solution (7.5mg/mL). At 10 days, we observed that most of the tails had fallen off, and the remaining 1-2 cm segment near the injection site was undergoing gradual necrosis. By 70 days, this segment had completely fallen off, and the wound had healed completely.

Application of CD to wounds created by cutting the tails of mice hastened wound healing compared to treatment with physiological saline, leading to complete wound closure approximately six days earlier (Supplementary Fig. S1A-B, E). In one group, prolonged exposure to CD led to significant damage to the excised wound and surrounding normal skin tissue. Reducing the exposure allowed the damaged skin to regenerate rapidly (Supplementary Fig. S1C). CD also facilitated the oxidation of cellular tissue debris near the wound (Supplementary Fig. S1F).

### Unilateral Tumor Experiments in C57BL/6 Mice

On the fourth day following the subcutaneous inoculation of LLC cancer cells in C57BL/6 mice, intratumoral injections with a lower dose of CD (8 mg/mL) were initiated, delivering 0.2 mL per mouse every other day, for a total of three doses. By the sixth day, these CD intratumoral injections significantly reduced tumor growth compared to the controls (Fig. 4A, Supplementary Fig. S2B). Tumors injected with CD also exhibited marked necrosis, indicating a strong necrotic effect on the tumor tissue (Supplementary Fig. S2A).

**Fig. 4.**
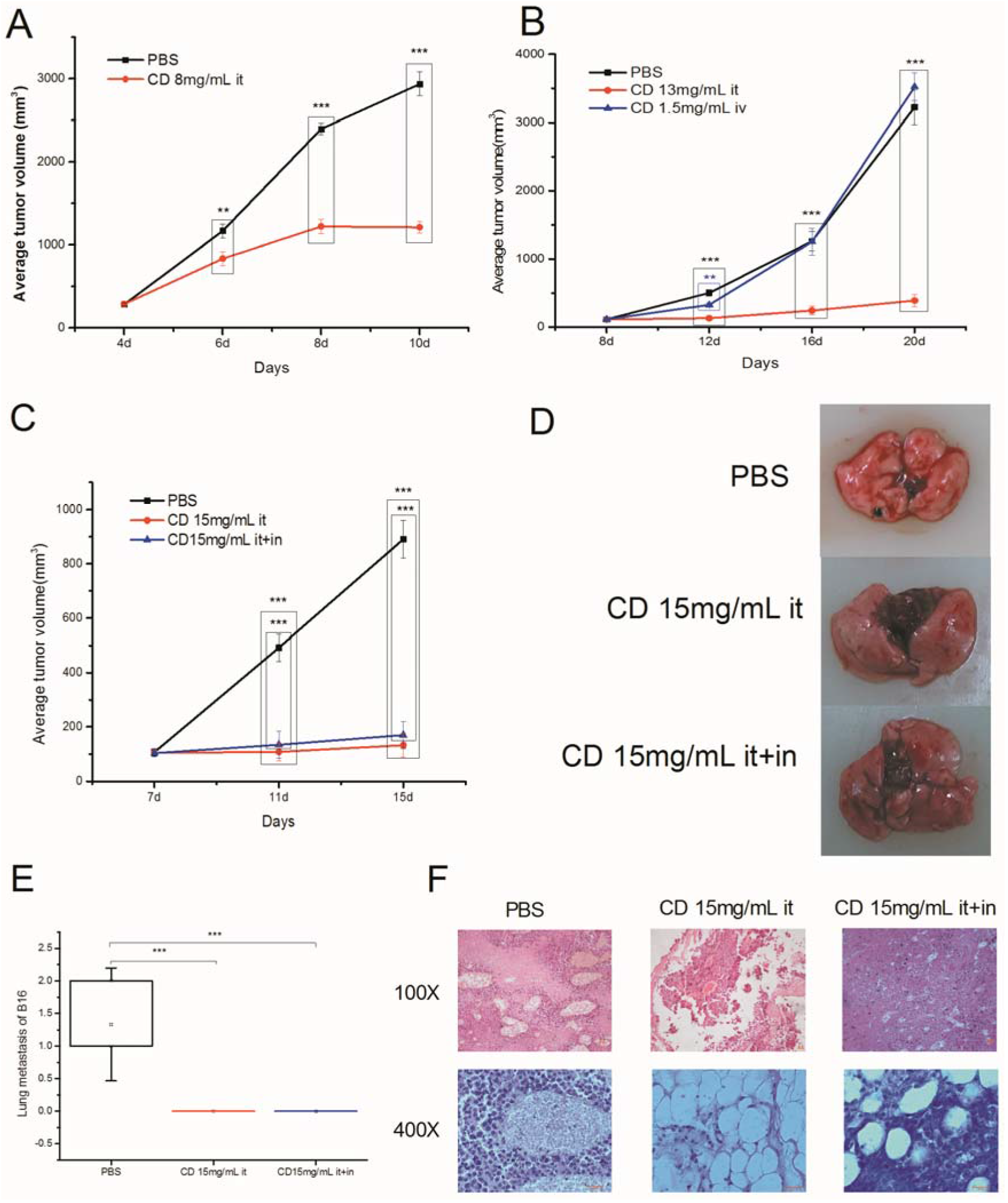
In vivo anti-cancer assay of CD through intratumoral injection in C57BL/6 mice. (A) Subcutaneous tumor volume of LLC cancer cells. One group received intratumoral injection of PBS as a control, while another group received intratumoral injection of CD (8mg/mL). (**P < 0.01, *** P < 0.001, two-tailed t-test; n = 10 mice per cohort). Error bars represent mean ± SEM. (B) Subcutaneous tumor volume of B16 cells, with one group receiving intratumoral injection of PBS as a control, another group receiving intratumoral injection of CD (13 mg/mL), and a third group receiving intravenous injection of CD (1.5 mg/mL). (**P < 0.01, *** P < 0.001, two-tailed t-test; n = 10 mice per cohort). Error bars represent mean ± SEM. (C) Tumor volume of subcutaneous and lung metastasis models established by subcutaneous and tail vein injection of B16 cells, with one group receiving intratumoral injection of PBS, another group receiving intratumoral injection of CD (15 mg/mL), and a third group receiving intratumoral injection of CD (15mg/mL) along with daily inhalation of CD gas. (*** P < 0.001, two-tailed t-test; n = 9 mice per cohort). Error bars represent mean ± SEM. (D) Lung metastasis nodules counted on day 21 in the subcutaneous and lung metastasis models. (E) Lung metastasis nodules counted on day 21. (*** P < 0.001, two-tailed t-test; n = 3 mice per cohort). Error bars represent mean ± SD. (F) Histological images of subcutaneous and lung metastasis models on day 17 stained with H&E (100x and 400x magnification).

To further investigate the therapeutic potential of CD, the dosage was increased to 13 mg/mL, administered at 0.2 mL every 4 days for a total of three doses. Additionally, a B16 tumor model was employed with an intravenous injection of CD at 1.5 mg/mL, given at 0.2 mL every 4 days for a total of five doses. Intratumoral injections achieved 74.1% tumor inhibition by day 4, whereas intravenous injections demonstrated a 34.8% inhibition (Fig. 4B). The intratumoral group consistently exhibited stronger tumor suppression, while the intravenous group did not maintain any inhibitory effect beyond the initial reduction in tumor burden.

On day 20, IL-1β and VEGF levels were assessed, and no significant differences were observed between the groups (Supplementary Fig. S3A, B). Furthermore, mice receiving intravenous injections of CD at 1.5 mg/mL showed no abnormal tail changes. In contrast, in previous non-tumor mouse trials, mice injected with 7.5 mg/mL displayed vascular damage, suggesting that higher concentrations of CD could adversely affect blood vessels.

In a B16 lung metastasis model, CD significantly inhibited tumor progression. Mice treated with daily inhalation of CD gas showed minimal effects on subcutaneous tumors (Fig. 4C). Upon examination on day 21, lung metastases were observed in control mice (1-2 foci per mouse, n=3), but no lung metastatic foci were found in mice receiving intratumoral CD injections or a combination of CD injection and inhalation (n=5, Fig. 4D, E).

Histological analysis performed on day 17 showed extensive necrosis and disruption of tumor architecture following CD injection (Fig. 4F). However, inhalation of CD gas caused lung damage without affecting the B16 lung metastases (Supplementary Fig. S4).

### Bilateral Tumor Experiments in BALB/c Mice

In a subsequent study, 4T1 tumor cells were injected into the mammary pad and axillary regions of BALB/c mice. Two groups of mice received varying concentrations of tumor cells: the small tumor group (ST) was given 4×10^5^ cells in the mammary pad and 2×10^5^ cells in the axillary region, while a large tumor group (LT) received 4×10^5^ cells in both regions. On the 15th day, CD (15 mg/mL) was administered exclusively into the mammary pad tumors of the experimental group. A total volume of 0.2 mL was injected every 3 days for a total of 6 doses.

We observed a significant reduction in tumor volume in the CD-treated groups, both in the large and small tumor groups. However, in the initial stages of treatment, the inhibition rate was notably higher in the small tumor group, particularly by the third day, achieving an inhibition rate of 72% (P < 0.01) compared to 30% (P < 0.10) in the large tumor group (Fig. 5A, B). The axillary tumors displayed a lower suppression rate due to the lack of direct effects from CD, and this reduction was not significant. It was only on days 21 and 27 that the CD-treated groups showed significantly higher inhibition rates compared to the control groups (Fig. 5C, D).

**Fig. 5.**
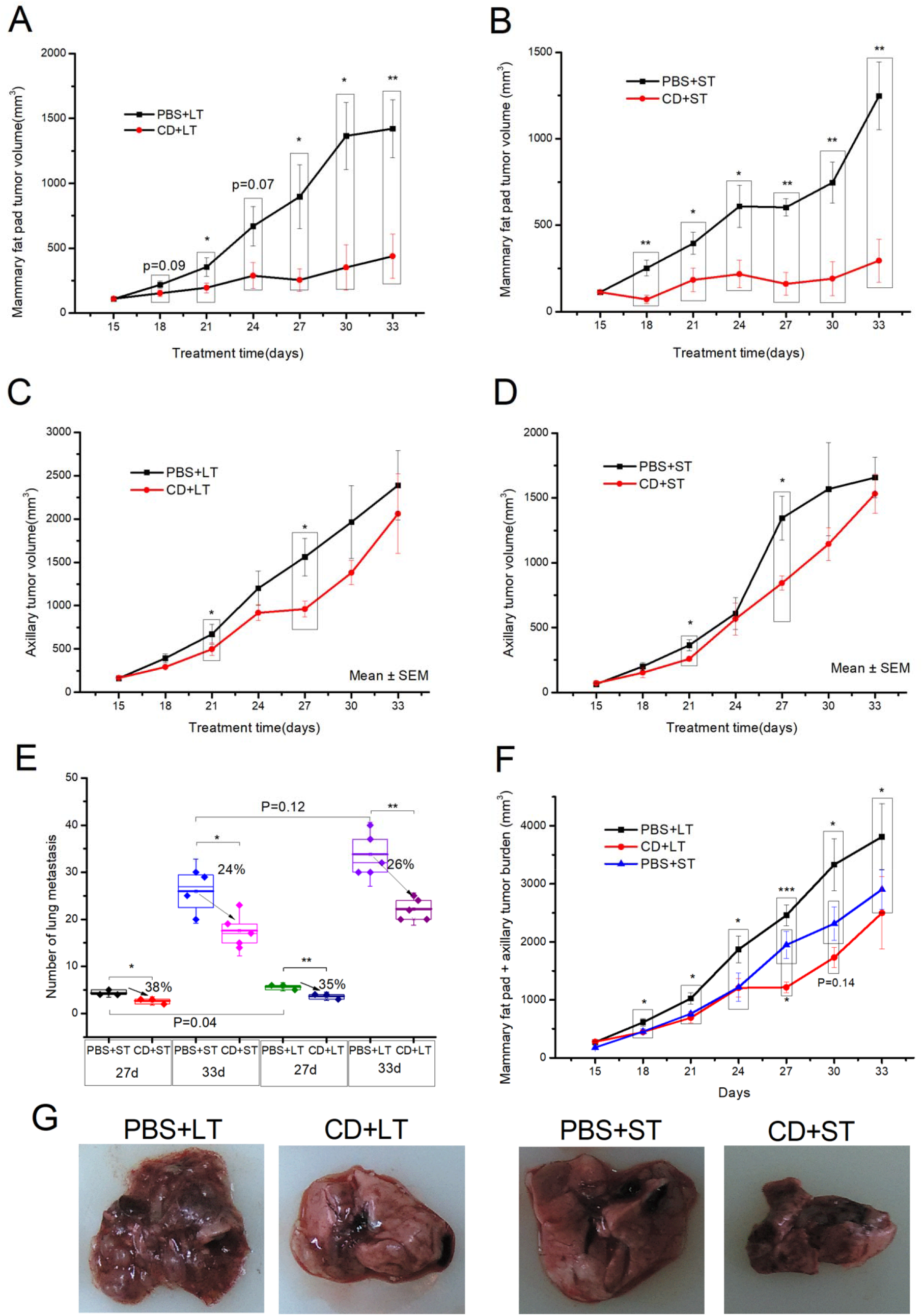
In vivo anti-cancer assay of CD through intratumoral injection in BALB/c mice. (A) Schematic representation of the subcutaneous and mammary pad injection of 4T1 tumor cells and the administration of CD. (B) Tumor volume of the large tumor group, with one group receiving intratumoral injection of PBS as a control (PBS+LT) and another group receiving intratumoral injection of CD (15 mg/mL) as a treatment (CD+LT). (*P < 0.05, ** P < 0.01, two-tailed t-test; n = 5-8 mice per cohort). Error bars represent mean ± SEM. (C) Tumor volume of the small tumor group, with one group receiving intratumoral injection of PBS as a control (PBS+ST) and another group receiving intratumoral injection of CD (15 mg/mL) as a treatment (CD+ST). (*P < 0.05, ** P < 0.01, two-tailed t-test; n = 4-8 mice per cohort). Error bars represent mean ± SEM. (D) Tumor volume of the large tumor group in the axilla (no injection). (*P < 0.05, two-tailed t-test; n = 5-8 mice per cohort). Error bars represent mean ± SEM. (E) Tumor volume of the small tumor group in the axilla (no injection). (*P < 0.05, two-tailed t-test; n = 4-8 mice per cohort). Error bars represent mean ± SEM. (F) Overall tumor burden, with the tumor volume of the large tumor group (mammary pad tumor injected with PBS) in the axilla added to represent the tumor burden of the control group, the tumor volume of the large tumor group (mammary pad tumor injected with CD) in the axilla added to represent the tumor burden of the treatment group, and the tumor volume of the small tumor group (mammary pad tumor injected with PBS) in the axilla added to represent the tumor burden of the control group. (*P < 0.05, ***P < 0.001, ANOVA, n = 5-8 mice per cohort). Error bars represent mean ± SEM. (G) Lung metastasis images on day 33, with the number of lung metastatic nodules roughly following the order: PBS+LT > CD+LT ≤ PBS+ST > CD+ST.

On days 27 and 33, lung metastasis was significantly reduced in the CD-treated groups compared to controls, with no significant delay differences between the small and large tumor groups (Fig. 5E, G). To assess overall tumor burden, we combined mammary pad and axillary tumor measurements. From day 24, the growth curve of the large tumor treatment group deviated significantly from that of the small tumor control group, indicating a notably lower growth rate (Fig. 5F). However, this immune enhancement was transient, as indicated by the convergence of growth curves after 30 days.

On day 27, no significant changes were observed in CD4+ T cells and NKp46 cells in the plasma and spleen, but there was an increase in CD11b+Gr-1+ myeloid cells in the treatment group compared to controls (Fig. 6A, B). The large tumor group exhibited a significant decrease in plasma CD8+ T cells, but an increase in the spleen, compared to the controls in the small tumor group.

**Fig. 6.**
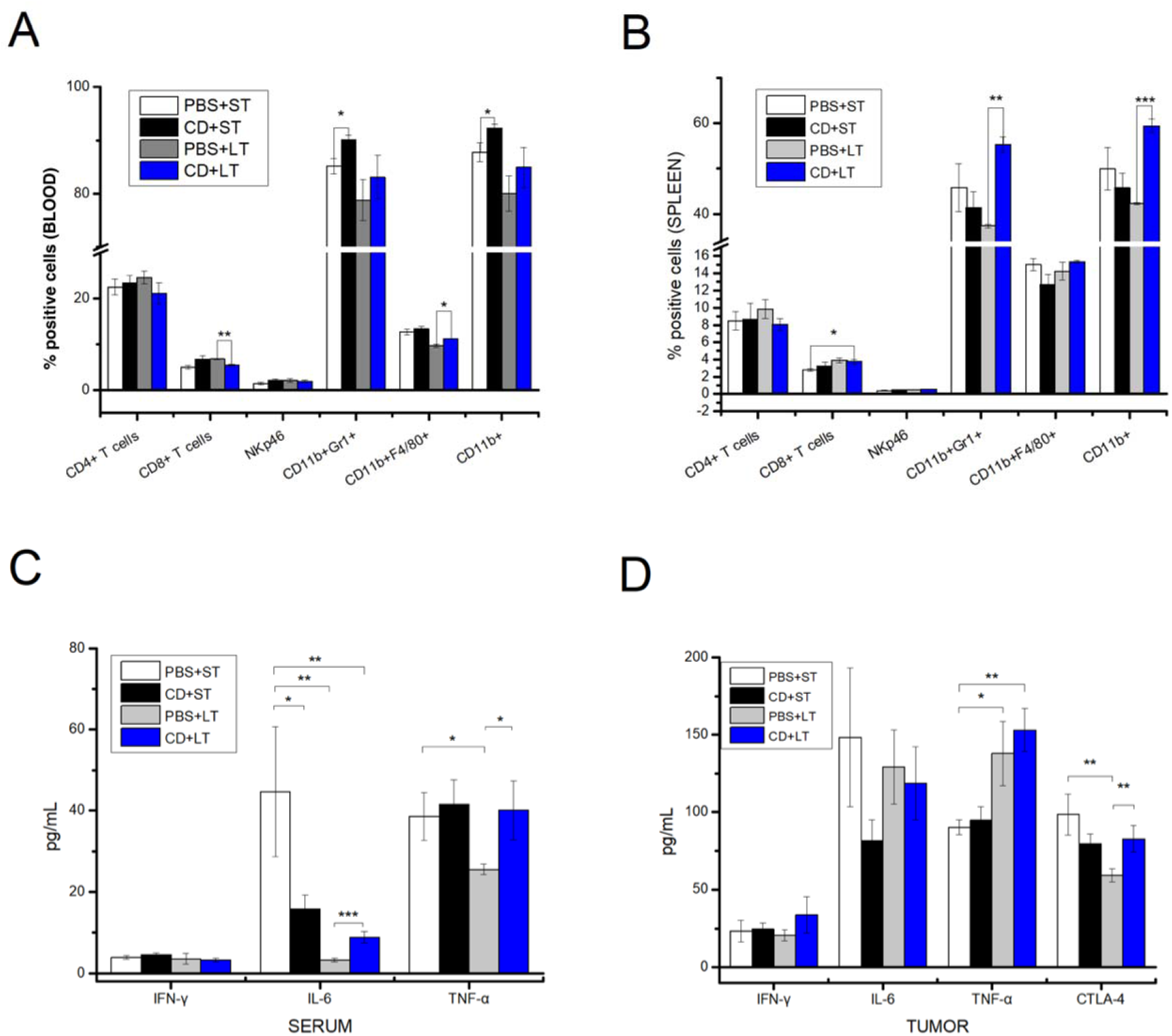
Comprehensive analysis of the therapeutic effects of intratumoral CD injections in BLAB/c mice inoculated with 4T1 cells. (A) Flow cytometric analysis of plasma samples from mice at day 27, demonstrating quantification of CD4, CD8, NKp46, CD11b+Gr-1+, CD11b+F4/80+, and CD11b+ positive cells. (B) Flow cytometric analysis of spleen samples from mice at day 27, showing quantification of CD4, CD8, NKp46, CD11b+Gr-1+, CD11b+F4/80+, and CD11b+ positive cells. (C) Cytokine analysis of serum samples from mice at day 33. (D) Cytokine analysis of tumor samples from mice at day 33. (A-D) Data are presented as the mean ± SEM. Differences between therapy groups were assessed using two-way ANOVA followed by Dunnett’s multiple comparisons tests (B-C: n=3; D-E: n=5).

By day 33, significant differences in cytokine levels were noted in serum and tumor tissues. TNF-α levels increased during later tumor development stages, higher in the large tumor group post-treatment compared to controls (Fig. 6C). Changes in TNF-α in tumor tissues mirrored those in serum, with IL-6 alterations suggesting interaction effects (Fig. 6D).

## Discussion

The cytotoxic effects of CD on both cancerous and normal cells are consistent with existing research, indicating that ROS can trigger both necrotic and apoptotic cell death [10]. This suggests that systemic administration of CD might cause widespread damage to normal tissues. However, the notable tissue damage observed after subcutaneous injection, although reversible, suggests that targeted delivery methods, such as intratumoral administration, might be more appropriate for the therapeutic application of CD. The reversible nature of CD-induced damage to normal tissues may allow for higher doses during intratumoral injections, potentially leading to more effective cancer cell elimination, even with some penetration into normal tissues. Additionally, CD’s ability to indiscriminately kill various cancer cell types without developing drug resistance underscores the advantage of direct intratumoral administration, ensuring effective and sustained cancer cell eradication.

The tail necrosis observed after tail vein injection raises concerns about CD’s potential to damage blood vessels, which may impact its use for intravenous applications. The eventual loss of the distal tail segment underscores the long-term ischemic damage that could result from systemic CD exposure. Conversely, when CD is administered via intratumoral injection, it might damage tumor vasculature, providing a potential therapeutic advantage despite the lack of selectivity.

Neutrophils are known to produce hypochlorous acid, an oxidizing agent similar to ROS, which has been shown to accelerate skin wound healing and promote skin rejuvenation [11]. Similarly, hydrogen peroxide, another ROS, has been found to facilitate sensory axon regeneration in zebrafish skin [12]. The accelerated wound healing observed in this study with CD supports these findings. However, the damage caused by prolonged exposure to CD highlights the need for careful regulation of exposure time and dosage to avoid adverse effects on healthy tissues. CD’s ability to oxidize cellular debris near wounds and promote tissue clearance may further enhance its role in regenerative processes (Supplementary Fig. S1F).

Ethanol ablation is a globally significant treatment modality because it requires no advanced technology beyond ultrasound. Ethanol is widely available, cost-effective, and the skills needed for its administration are easily taught and learned. Additionally, the procedure can be repeated with only local anesthesia. However, while generally considered safer, ethanol ablation appears to be less effective than surgical interventions [13]. This reduced efficacy may be due to the longer retention time required within tumors for effective cancer cell eradication.

Injecting CD into tumors could offer similar advantages to ethanol ablation. Due to CD’s rapid oxidative properties, it may swiftly kill cancer cells upon penetration without the need for prolonged retention. Thus, achieving the desired therapeutic effects might be possible by primarily increasing the effective dosage of CD.

Upon examining the wound shapes formed, we observed that the spread of CD was uniform around the injection site, with no distinct layers forming in the center. The entire elliptical wound exhibited a consistent intensity of damage. These observations suggest that increasing the dose or concentration of CD during injection could allow a single treatment to destroy a larger portion of the tumor, enhancing cancer cell necrosis. In specific scenarios, CD might penetrate from the tumor into surrounding healthy tissue, ensuring comprehensive tumor destruction while keeping complications from damage to normal tissue within acceptable limits.

Intratumoral injection of CD at lower doses (8 mg/mL) demonstrated significant tumor suppression by day 6, and the necrosis observed in CD-treated tumors suggests that CD effectively induces tumor cell death, potentially through both apoptotic and necrotic mechanisms (Supplementary Fig. S2A). This aligns with prior studies indicating that ROS can induce both necrosis and apoptosis [10], independent of the cell type.

When the CD dose was escalated to 13 mg/mL in the B16 tumor model, a marked 74.1% tumor suppression was observed by day 4, compared to the 34.8% inhibition observed with intravenous injections. This discrepancy may stem from the rapid oxidative effects of CD when directly injected into the tumor, allowing it to act more efficiently. In contrast, intravenous administration seems to have only a transient effect on tumor size, potentially due to systemic clearance or limited tumor penetration.

By comparing CD injections into the tail veins of non-tumor-bearing and tumor-bearing mice, we observed that higher concentrations of CD (7.5 mg/mL compared to 1.5 mg/mL) result in significant vascular damage, suggesting that CD might disrupt tumor vasculature as well. This effect could enhance its therapeutic efficacy when administered directly into tumors. Due to poorly regulated angiogenesis, tumor-associated blood vessels often exhibit defects and leaks [14]. We hypothesize that higher concentrations of CD could more readily penetrate these compromised vessels, causing additional damage to both tumor cells and their vasculature. This phenomenon may contribute to CD’s enhanced tumor-killing effect in areas otherwise inaccessible to the CD solution.

The combined induction of apoptosis and necrosis by CD is particularly relevant given that both forms of cell death have been shown to trigger anti-tumor immune responses [15, 16]. In our B16 lung metastasis model, intratumoral injection of CD not only suppressed primary tumor growth but also significantly inhibited metastasis, likely due to the activation of the body’s anti-tumor immune response. Interestingly, while CD inhalation caused some lung damage (Supplementary Fig. S4), it did not influence lung metastasis, suggesting that CD’s systemic effects on immune modulation are primarily driven by direct tumor injection rather than inhalation.

Histological analysis further supports these findings, with extensive necrotic centers observed in CD-injected tumors, indicating substantial tumor destruction. This, combined with the absence of lung metastases in CD-treated mice (Fig. 4D), demonstrates that CD may offer a dual mechanism of action by directly ablating the tumor and simultaneously activating systemic immune responses against tumor cells.

We observed consistency between the results of the bilateral tumor experiments in BALB/c mice and the unilateral tumor experiments in C57BL/6 mice, where intratumoral injection of CD significantly inhibited tumor growth in the treatment group. Moreover, there was a proportional relationship between the CD dosage and inhibition rate relative to tumor size. Additionally, tumors on the contralateral side that were not injected also exhibited some degree of inhibition, suggesting a possible immune suppression effect, although it appears to be only transient.

The presence of a primary tumor has been shown to decrease immunocompetence, but this can be improved by surgical resection of the primary tumor [17]. In our study, we investigated the effects of intratumoral CD injection for tumor ablation on the growth trajectory of tumors in different sizes. We hypothesized that if intratumoral CD injection can enhance the antitumor immune response, it would result in a significantly lower growth curve in the large tumor treatment group compared to the untreated growth curve of the small tumor group.

Our results showed that from day 24, the growth curve of the overall tumor burden in the large tumor treatment group deviated from that of the small tumor control group, with a notably lower growth rate. This suggests that intratumoral CD injection not only directly ablates the tumor and restores the dominant immune capability but also provides additional immune enhancement. However, it is important to note that the immune enhancement observed was transient, as the growth curves of both groups started to converge after 30 days. These findings indicate that intratumoral CD injection has the potential to enhance the immune response against tumors, but further studies are needed to understand the duration and sustainability of this immune enhancement.

On day 27, no significant changes were observed in CD4+ T cells and NKp46 cells in the plasma and spleen. However, the treatment group showed a significant increase in CD11b+Gr-1+ myeloid cells compared to the control group. In the large tumor group, the treatment group exhibited a significant decrease in CD8+ T cells in the plasma compared to the control group. Conversely, in the spleen, the treatment group in the large tumor group demonstrated a significant increase in CD8+ T cells compared to the control group in the small tumor group. These findings are consistent with the enhanced immune response induced by intratumoral CD injection. It is worth noting that CD11b+Gr-1+ cells possess immunosuppressive properties and contribute to the promotion of cancer metastasis [18], which may explain the transient nature of the immune response stimulated by intratumoral CD injection.

On day 33, cytokine evaluations revealed significant differences in TNF-α and CTLA-4 levels, which reflect a complex dynamic of immune activation and suppression. These changes indicate an ongoing but reduced antitumor immune response after the initial CD injection. Prolonging immune activation and overcoming immunosuppression, possibly through combination therapies or targeted immune pathway strategies, could enhance the efficacy and duration of CD injections as a cancer treatment. T-cell exhaustion has been identified as a possible mechanism for transient effects [19]. Further investigation is essential to elucidate factors affecting response persistence and optimize treatment protocols.

Our study highlights CD’s properties similar to ROS, such as promoting tissue regeneration and performing comparable functions. Given the significant role of injury-induced ROS production in tissue regeneration [20], we’ve confirmed that CD can facilitate tissue regeneration in injured areas, akin to ROS. Tumors can be regarded as non-healing wounds [21], presenting an opportunity to leverage CD’s regenerative capabilities in cancer treatment to expedite wound healing and improve patient outcomes. Direct intratumoral administration of CD conveniently and sustainably kills cancer cells without the risk of developing drug resistance. Compared to conventional cancer treatments, CD offers numerous advantages, significantly reducing disease complications and associated risks, thus providing a safer and more effective alternative. While higher CD concentrations may pose a risk of vascular damage, they also offer potential benefits in tumor ablation.

A notable limitation of this study is the absence of direct measurements of the potential destructive effects of high-concentration CD within tumors, warranting further exploration. Additionally, our research lacks injection experiments involving larger tumors, which could impede a comprehensive understanding of the penetration patterns of CD solutions within tumor structures. Future studies should include a wider range of tumor sizes to accurately assess CD’s penetration capabilities and therapeutic efficacy across different tumor volumes.

We propose the concept of intratumoral CD injection to facilitate direct tumor ablation without encountering resistance, while concurrently enhancing antitumor immunity. This approach minimizes damage to healthy tissues and leverages regenerative potential post-ablation, optimizing cancer treatment outcomes. Observations from bilateral tumor models suggest that CD may stimulate an antitumor immune response, indicating a need for more in-depth research to better understand the underlying mechanisms.

As immune checkpoint inhibitors are revolutionizing oncology, numerous clinical trials are evaluating strategies like intratumoral delivery and tumor tissue-targeted compounds to enhance local bioavailability and increase the effectiveness of immunotherapies [22]. Technological advancements position intratumoral CD administration as a promising, efficient, and patient-friendly cancer treatment strategy, potentially extending patient survival and reducing treatment burden. This strategy could redefine cancer management, likening it to treating chronic conditions.

To realize this potential, further research is crucial to refine CD administration protocols and dosages, investigate synergistic effects with other therapies, and understand its regenerative and ablative effects on tumor tissue, as well as its capability to elicit a systemic antitumor immune response. These findings lay a crucial foundation for continued exploration of CD in oncological applications. The ongoing exploration of CD highlights its prospective value as a transformative tool in cancer therapy, signaling new opportunities for enhancing patient care and clinical outcomes.

## Author contributions

Xuewu Liu contributed to the study design and development of the research framework. Zhaoyang Liu, Xuewu Liu, Jiao Zhang, and Xueyan Liu were responsible for conducting the experiments. Xuewu Liu, Shuangning Liu, and Jiao Zhang performed data analysis and contributed to the writing of the manuscript. Shuangning Liu created the main figures. All authors have reviewed and approved the final version of the manuscript for publication.

## Data Availability Statement

All relevant data are available upon request from the corresponding author.

## Abbreviations

SOD: Superoxide dismutase
ROS: Reactive Oxygen Species
PDT: Photodynamic therapy
CD: Chlorine dioxide
DAMPs: Death-associated molecular patterns
HCMs: Human cardiac myocytes
HUVECs: Human vascular endothelial cells
NSCLC: non-small cell lung cancer
LT: Large tumor
ST: Small tumor

