## Appendix A. Supplementary Figure for "Intratumoral Delivery of Chlorine Dioxide Exploits its ROS-like Properties: A Novel Paradigm for Effective Cancer Therapy"

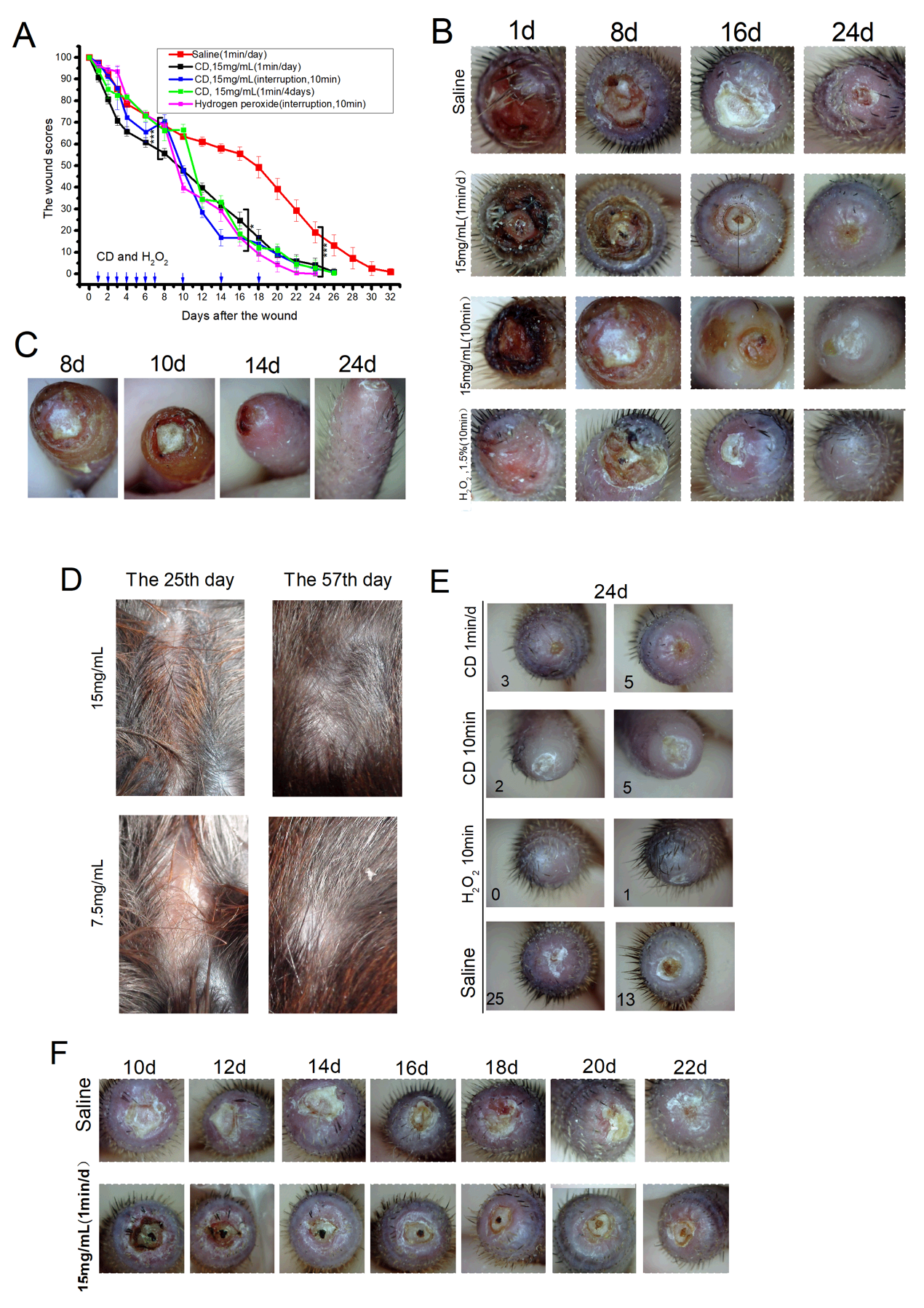

**Fig S1. Effects of CD on tissue damage and wound healing in normal tissues.** (A) Tail injury scores (wound healing) in C57BL/6 mice. One group was treated with 15 mg/mL CD for 1 minute daily, another group was treated with saline for 1 minute daily, the third and fourth groups were treated with 15 mg/mL CD and 1.5% hydrogen peroxide for 10 minutes daily from day 1 to day 7, with three additional treatments on day 10, 14, and 18 for 10 minutes each; the fifth group was treated with 15 mg/mL CD for 10 minutes on day 1, 4, 8, 12, 16, and 20. (*** P < 0.001, Mann-Whitney test; n = 4 mice per cohort). Error bars represent mean ± SD. Prolonged treatment with CD on the wound caused direct damage to the surrounding tissue, resulting in fluctuation in both damage and repair. Similar effects were observed with hydrogen peroxide treatment. (B) Photographs of the tail wounds at four time points, showing the process of wound healing. One group was treated with 15 mg/mL CD for 1 minute daily, another group was treated with saline for 1 minute daily, the third and fourth groups were treated with 15 mg/mL CD and 1.5% hydrogen peroxide for 10 minutes daily from day 1 to day 7, with three additional treatments on day 10, 14, and 18 for 10 minutes each. (C) Significant tissue damage was observed in the normal tissue near the tail wound when immersed in a 15 mg/mL CD solution for 10 minutes daily from day 1 to day 7, 10, 14, and 18. However, when the continuous treatment was stopped, the created wounds and damaged tissue healed and regenerated almost synchronously. (D) Photographs of C57BL/6 mice injected with 15 mg/mL and 7.5 mg/mL CD solutions subcutaneously on the back on day 25 and day 57, showing the initiation of new hair growth around the injection site. (E) Photographs of tail wounds in two mice treated with CD (immersed in a 15 mg/mL CD solution for 1 minute daily) and two mice treated with saline after 24 days of tail injury. Photographs of tail wounds in each group of two mice treated with CD and hydrogen peroxide (immersed in a 15 mg/mL CD solution for 10 minutes and a 1.5% hydrogen peroxide solution for 10 minutes) on day 1 to day 7, 10, 14, and 18 are shown, with the lower numbers indicating the wound scores. (F) Representative photographs of tail wounds in C57BL/6 mice treated with saline and CD from day 10 to day 22. The presence of accumulated debris attached to the wound indicates that CD treatment can clear these deposits.

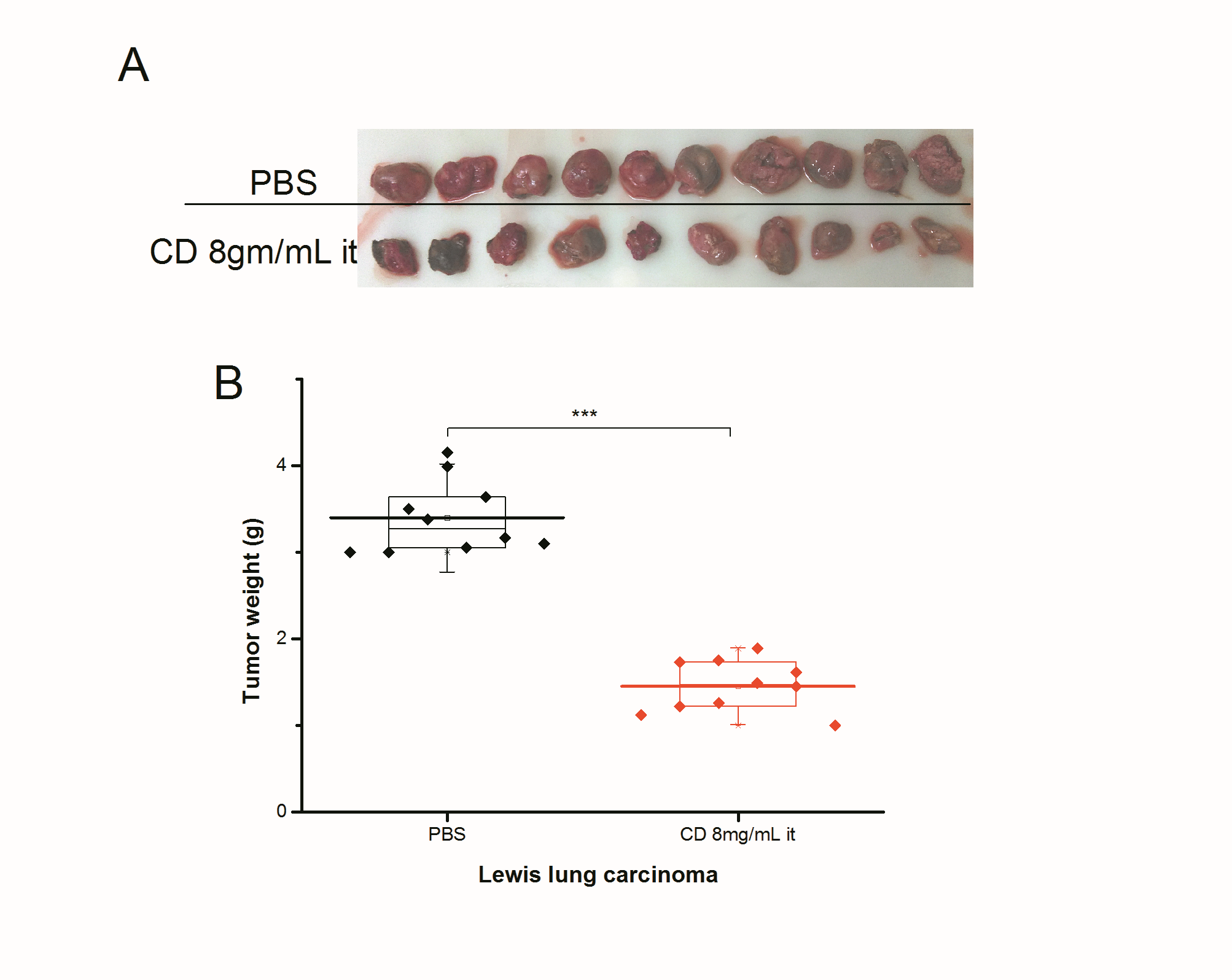

**Fig S2. Inhibition of Lewis lung carcinoma (LLC) subcutaneous tumor growth by CD treatment.** (A)Tumors were measured on day 10 after inoculation. One group received intratumoral injections of PBS as a control, while another group received intratumoral injections of CD (8 mg/mL). (B) Tumor weight data. (*** P < 0.001, two-tailed t-test; n = 10 mice per cohort). Error bars represent mean ± SD.

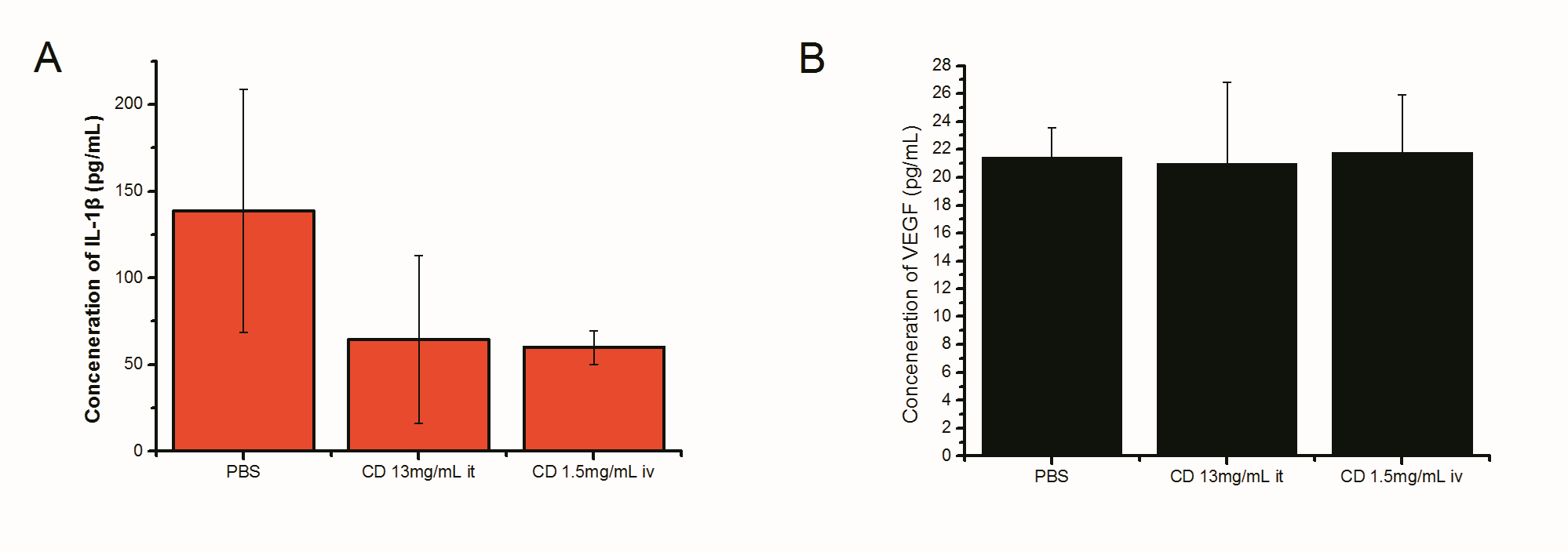

**Fig S3. Cytokine detection in C57BL/6 mice subcutaneously inoculated with B16 cells.** One group received intratumoral injections of PBS as a control, another group received intratumoral injections of CD (13 mg/mL), and a third group received intravenous injections of CD (1.5 mg/mL). On day 20, the levels of (A) IL-1β (n=3-5), Error bars represent mean ± SD, and (B) VEGF (n=5), Error bars represent mean ± SD, in the tumor cells were measured.

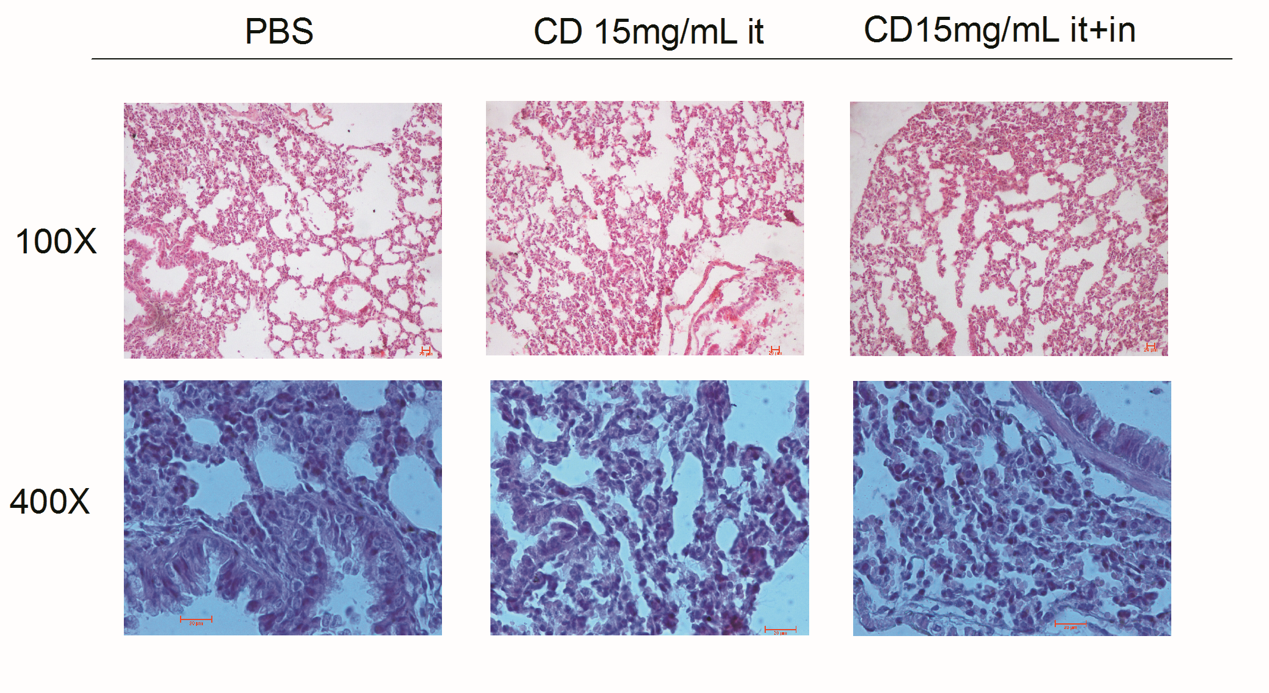

**Fig. S4. C57BL/6 mouse B16 subcutaneous and lung metastasis model.** The groups consisted of a control group (injected with PBS), a group receiving intratumoral injections of CD (15 mg/mL it), and a group receiving intratumoral injections plus daily inhalation of CD gas (CD 15 mg/mL it+in). On day 17, lung tissues were stained with H&E and examined at 100x and 400x magnification.
